# OsAAI1-OsMADS25 module orchestrates root morphogenesis by fine-tuning IAA in drought stressed rice

**DOI:** 10.1101/2024.06.03.597090

**Authors:** Ning Xu, Rui Luo, Qing Long, Jianmin Man, Jiaxi Yin, Haimin Liao, Meng Jiang

**Author notes:** These authors contributed equally to this work (Ning Xu, Rui Luo).

## Abstract

Indole-3-acetic acid (IAA) plays a critical role as a plant hormone in regulating the growth and development of the root system in plants, particularly in enhancing their ability to withstand abiotic stress. In this study, we found that overexpression of *OsAAI1* promoted the growth of rice root system. The length of primary root, the number of lateral roots, the density of lateral roots, and the number of adventitious roots of overexpression of *OsAAI1* (OE19) were significantly better than those of the wild type (ZH11) and the mutant line (*osaai1*), and the IAA content of OE19 was significantly higher than those of ZH11 and *osaai1*. We also found that exogenous application of IAA could compensate for the root growth defect caused by the *osaai1* mutation. OE19 had the highest number and widest distribution of total roots under the water-cut drought treatment, and exogenous application of IAA attenuated the growth inhibitory effect of drought stress on *osaai1*. Our study also revealed that OsAAI1 interacts with the MADS-box family transcription factor OsMASD25. Additionally, the application of IAA helped alleviate the growth inhibitory effects of drought stress on *osmads25*.Importantly, OsMADS25 interaction with OsAAI1 was found to enhance the transcriptional expression of its downstream target genes *LAX1* and *OsBAG4*, which are crucial genes in rice’s response to drought stress. These findings suggest that OsAAI1 and OsMADS25 are crucial in rice’s drought acclimation process by regulating downstream gene expression and influencing the IAA signaling pathway.

**Author summary:** The root system is a crucial organ for crop plants as it facilitates the absorption of water and nutrients, contributing to their drought resistance. Indole-3-acetic acid (IAA) plays a pivotal role in the growth of various types of roots in plants. Under drought stress conditions, changes in IAA levels and transport can impact the morphology of plant roots. This research illustrates that OsAAI1 positively influences rice root development and enhances the plant’s response to drought stress through the auxin signaling pathway. The study reveals a physical interaction between OsAAI1 and the transcription factor OsMADS25. This interaction boosts the expression of the auxin synthesis gene *OsYUC4* and suppresses the auxin inhibitory factor *OsIAA14*, thereby promoting the auxin signaling pathway, stimulating rice root growth, and enhancing the plant’s ability to withstand drought. Furthermore, the interaction between OsAAI1 and OsMADS25 has been found to also positively affect the expression of the genes *LAX1* and *OsBAG4*, which is associated with activated drought resistance in rice plants.

## Introduction

Roots are nutrient organs in the underground part of the plant with the functions of absorption, fixation, transport, synthesis, storage, multiplication and secretion. Its main function is to absorb water and inorganic salts in the soil, and to transport the absorbed substances to the above ground for plant growth[1]. Rice roots can be divided into primary roots, lateral roots and adventitious roots[2]. Primary roots and lateral roots are developed from the radicle, lateral roots begin with a few pericytes, which take on the characteristics of the initiating cell, and then divide asymmetrically to produce a lateral root primordium (LRP), which continues to grow and eventually emerges from the primary root[3,4]. Adventitious roots are roots of uniform thickness produced by different parts of the radicle, stem, leaves or old roots[5]. Multiple research projects have demonstrated the significant importance of rice root growth and development in managing drought conditions[6–8]. Three primary mechanisms of adaptation have been observed in rice under drought conditions: (i) adjustments in osmotic levels in roots when there is limited soil moisture; (ii) heightened root penetration into the soil; and (iii) enhanced root depth, density, and root-to-shoot ratio when soil moisture levels are high[8]. Many studies have shown that rice plants with deeper roots exhibit greater tolerance to water scarcity[1].

Being one of the extensively researched plant radicals, Auxin participates in nearly all plant growth and developmental processes[9]. Indole-3-acetic acid (IAA) serves as the main auxin in the majority of plants, influencing root system structure and different root growth phases[10]. Current research indicates that auxin is crucial for enhancing plants’ tolerance to different abiotic stresses[11–13]. When plants are subjected to abiotic stresses, growth hormone homeostasis is disturbed, and IAA can potentially promote root growth by regulating genes involved in auxin biosynthesis, efflux, and conjugation and degradation, thereby resisting adversity stresses[12,14]. As stress-responsive substances, ROS are composed of hydrogen peroxide (H2O2), superoxide anion (O2 ^-^) and hydroxyl radical (OH^-^)[15]. Homeostasis of reactive oxygen species (ROS) in plant cells is crucial for cellular function, achieved through a careful balance between ROS production and scavenging mechanisms[16]. During periods of abiotic stress, such as exposure to environmental factors like high salinity or extreme temperatures, plants often accumulate high levels of ROS [17]. To counteract the potential damage caused by ROS, plants employ an enzyme scavenging system that relies on the actions of enzymes like superoxide dismutase (SOD), catalase (CAT), ascorbate peroxidase (APX), and glutathione peroxidase (GPX) to eliminate ROS from the cellular environment [16,18]. Numerous studies have demonstrated the role of the plant hormone indole-3-acetic acid (IAA) in promoting plant resistance to abiotic stress by enhancing ROS scavenging[19,20]. For instance, Gong et al. conducted experiments on cucumbers and found that the application of exogenous IAA led to an increase in IAA concentration in the leaves, resulting in elevated CAT activity that helped alleviate sodic alkaline stress[19]. Ma et al. similarly showed that the exogenous application of IAA could mitigate the negative effects of alkaline stress on plant growth by regulating the plant’s ROS scavenging system, promoting root development, and affecting the expression of genes involved in IAA biosynthesis, transport and catabolism[20]. These findings highlight the importance of IAA in enabling plants to cope with and adapt to challenging environmental conditions by modulating ROS levels and enhancing stress tolerance[21].

Alpha-Amylase Inhibitors (AAI), Lipid Transfer (LT), and Seed Storage (SS) Protein family (AAI_LTSS Protein family) is a distinctive set of proteins discovered in higher plants, consisting of 5 individuals. These individuals are referred to as Alpha-Amylase Inhibitors (AAIs) and the Seed Storage (SS) Protein subgroup (AAI_SS), the Hydrophobic Protein from Soybean (HPS)-like subgroup, the Non-specific lipid-transfer protein type 2 (nsLTP2) subgroup, the Non-specific lipid-transfer protein type 1 (nsLTP1) subgroup, and the Non-specific lipid-transfer protein (nsLTP)-like subgroup, respectively. Limited information is available concerning their involvement in controlling plant root growth. OsAAI1 is an AAI gene from the HPS-like subgroup and codes for three domains: the LTP2 domain, the hydrophobic seed domain, and the trypsin alpha amylase domain[21]. Our previous research demonstrated that increased levels of *OsAAI1* enhanced rice’s ability to withstand drought by activating both the ABA signaling pathway and the ROS scavenging pathway[21]. In this study, we further found that overexpression of *OsAAI1*(OE19) grew significantly better than the wild type Zhonghua 11 (ZH11) and mutant line (*osaai1*) in terms of primary root length, lateral root number, lateral root density, total root length, and number of adventitious roots. Additionally, the IAA concentration in OE19 was notably higher than ZH11 and *osaai1*, and that exogenous application of 10nM IAA compensated for the root growth defects induced by the *osaai1* mutation. In addition, we found that exogenously applied IAA could alleviate the inhibitory effect of drought stress on *osaai1*. In this study, we demonstrated that OsAAI1 interacts with OsMADS25 and that the exogenous application of IAA also alleviates the inhibitory effect of drought stress on *osmads25*. This study aimed to investigate how OsAAI1 and OsMADS25 constitute a regulatory network through interaction in response to drought stress and regulate ROS homeostasis and IAA-mediated root morphogenesis under drought stress in rice. The results suggest that OsAAI1 and OsMADS25 play an important role in drought acclimation in rice by interacting with each other, regulating the expression of downstream genes and ultimately influencing ROS homeostasis and IAA signaling pathways.

## Results

### *OsAAI1* regulates the development of primary root length and the formation of lateral root primordia in rice

Our phenotypic observations and data analysis of wild-type and transgenic lines grown normally for 14 days, the results showed that compared to ZH11, OE19 increased rice primary root length, lateral root number, lateral root density, adventitious root number and total root length, while the opposite was true for *osaai1* (Fig1A-F). Lateral root primordia staining was used to explore the effect of *OsAAI1* on the formation of lateral root primordia on rice primary roots, and it was found that the number and density of lateral root primordia on the primary roots of *osaai1* were significantly lower than those of ZH11, while the number and density of lateral root primordia of OE19 were significantly higher than those of ZH11(Fig1G, H), indicating that *OsAAI1* affected the initiation of lateral root primordia in rice. We made paraffin sections of the wild-type and transgenic lines at 4 days of growth to observe the differences between the cells of each line. The transverse section of the roots showed that OE19 had more thin-walled cells in the xylem and phloem than ZH11 and they were closely arranged. Moreover, OE19 was the first to show the lateral root primordia, and the longitudinal section of the roots showed that the cells in the elongation zone of OE19 were longer (Fig1I and S1FigA, B). We observed the root growth of wild-type and transgenic lines grown normally for 1–3 months in rhizotron and found that the root growth of OE19 was consistently the best, followed by ZH11, and *osaai1* was the worst (Fig2A-G, S2Fig and Table 1). Our previous study showed that OE19 had better root length and root weight than ZH11 and *osaai1* at the maturity stage[21], indicating that *OsAAI1* affected the root development of rice throughout the growth period. In summary, *OsAAI1* regulates primary root elongation by regulating the length of rice primary root cells.

**Table1.**
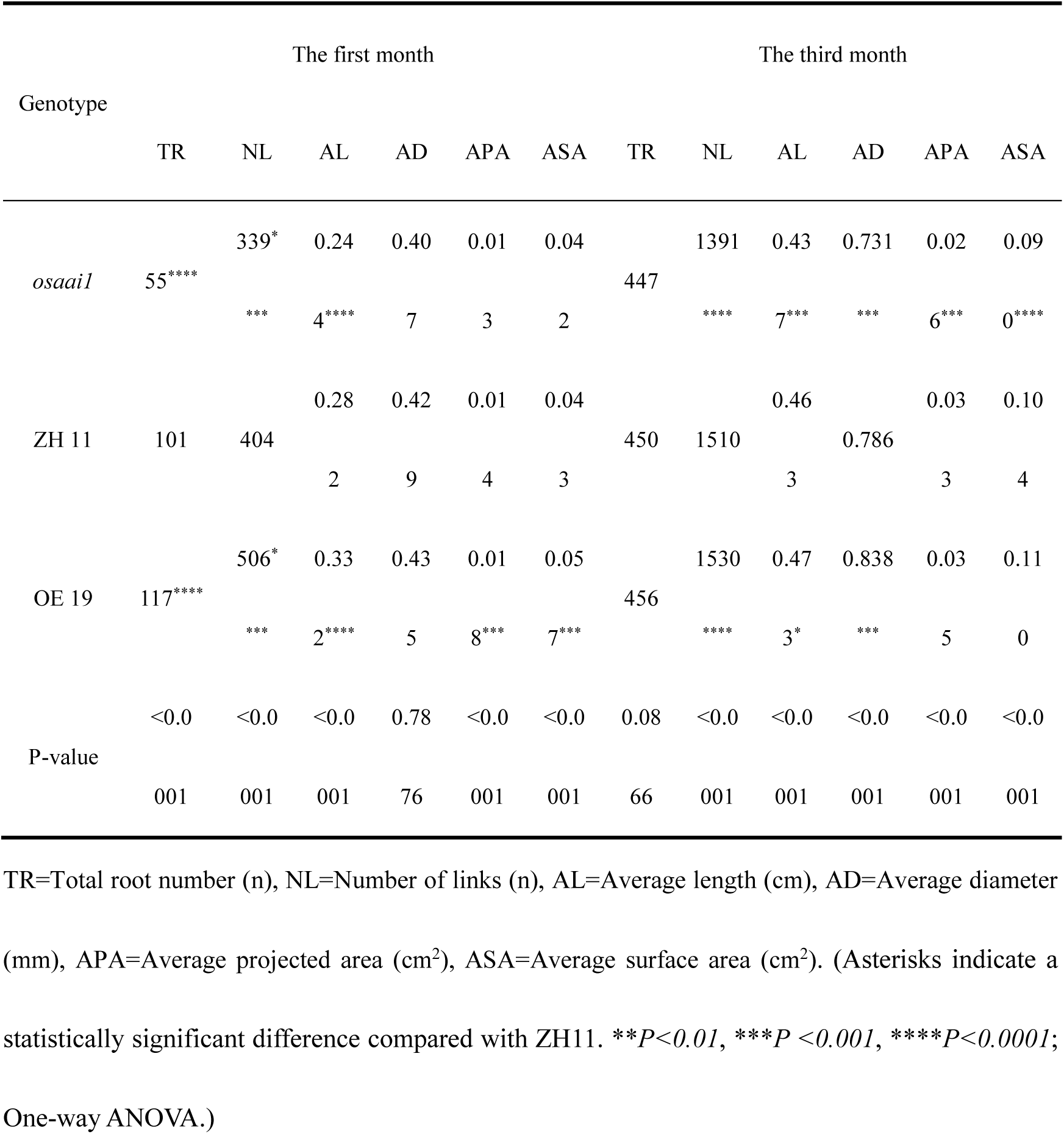
Statistical analysis of traits in wild-type and transgenic lines during the first and third months of normal growth.

**Fig 1.**
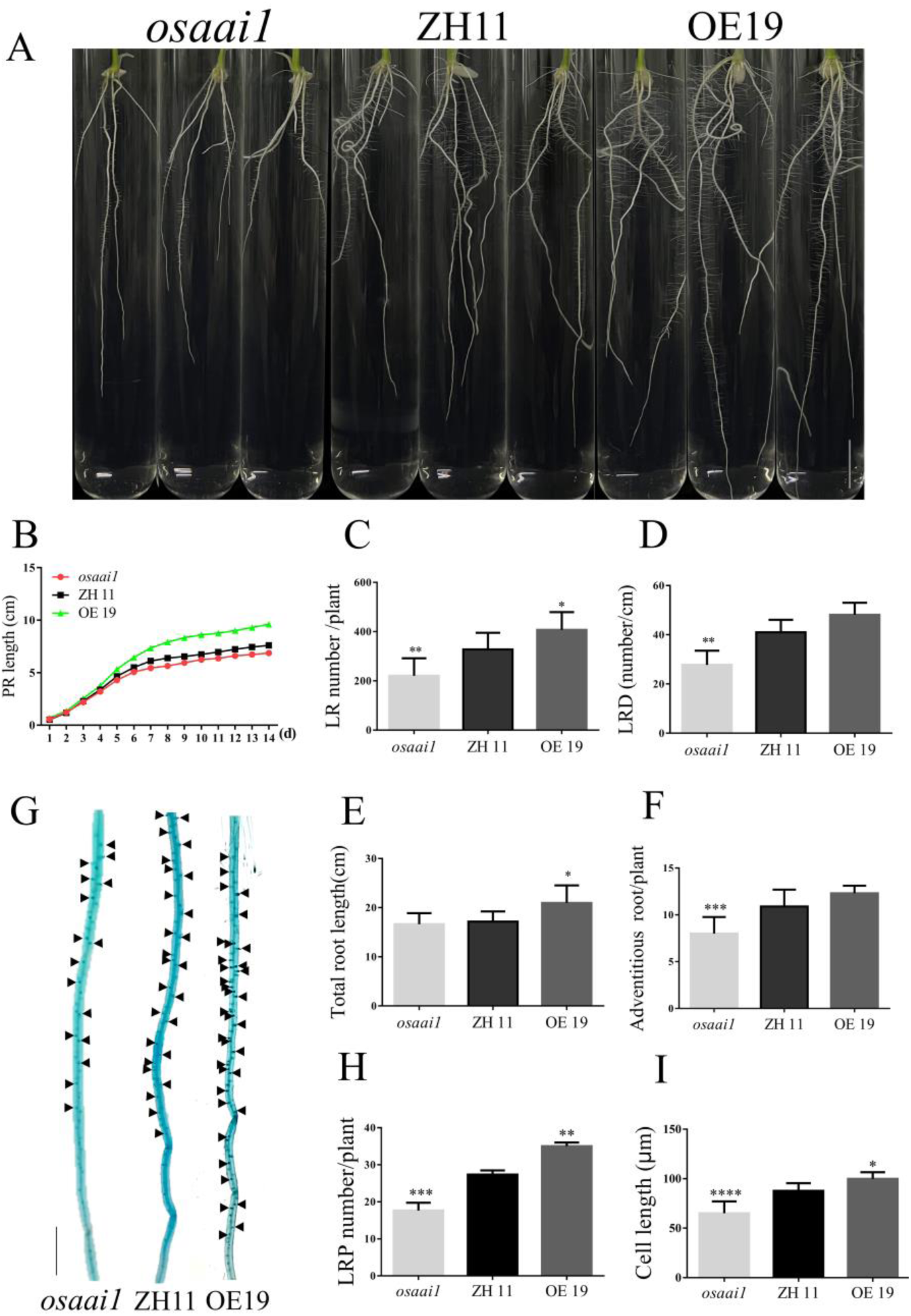
*OsAAI1* regulates the development of primary root length and the formation of lateral root primordia. Phenotype(A), scale bar = 1.5cm, primary root length(B), number of lateral roots(C), lateral root density(D), total root length(E), and number of adventitious roots(F) of wild type and transgenic lines at 14 days of normal growth. Lateral root primordia staining(G), bar = 0.5cm, lateral root primordia density(H) of wild type and transgenic lines at 5 days of normal growth. Cell length statistics of paraffin sections(I) of wild-type and transgenic lines at 5 days of normal growth. (Asterisks indicate a statistically significant difference compared with ZH11. \*\**P<0.01*, \*\*\**P <0.001*, \*\*\*\**P<0.0001*; One-way ANOVA.)

**Fig 2.**
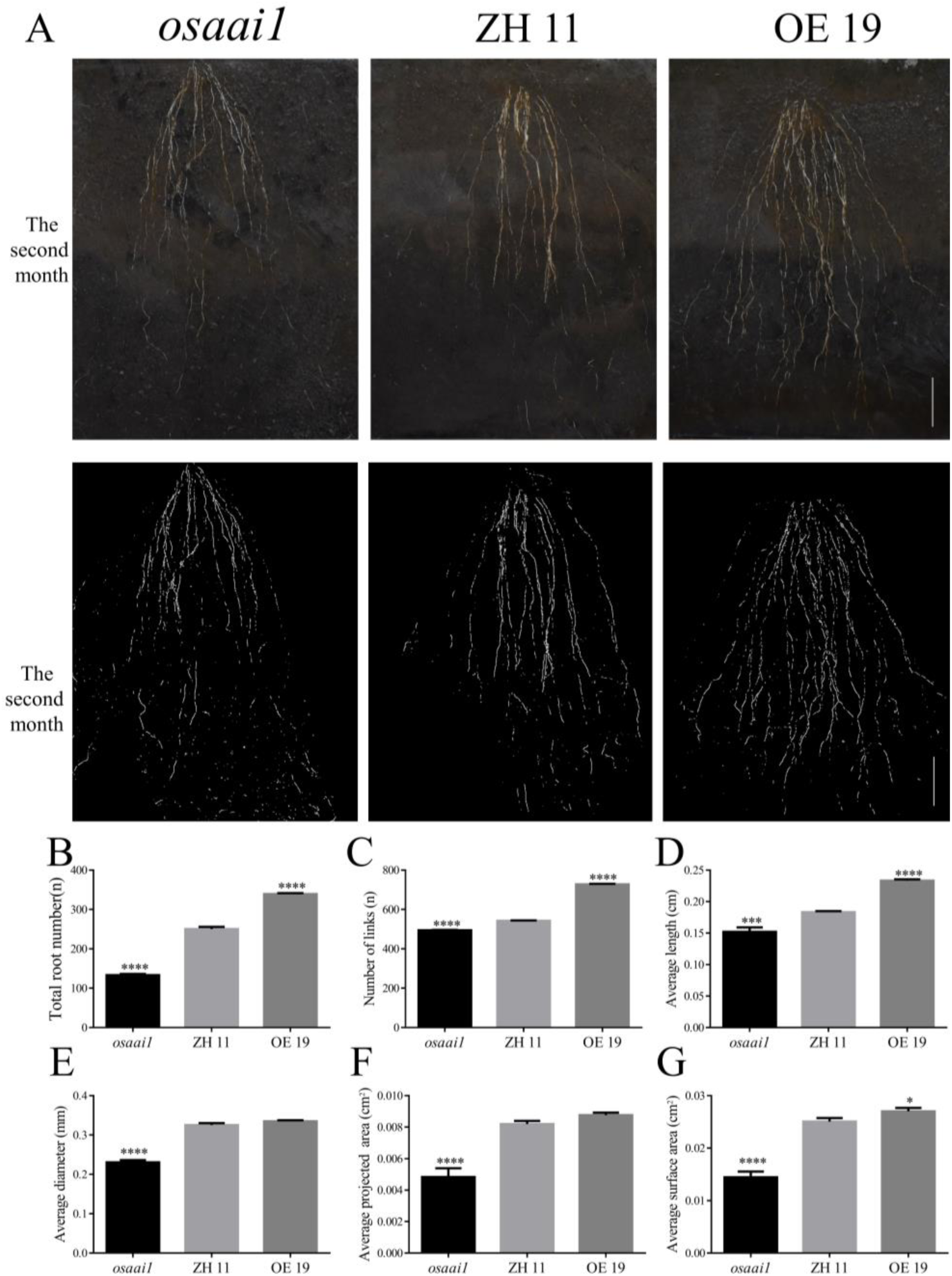
Root system architecture (RSA) of wild type and transgenic lines grown in rhizotrons. RSA of wild type and transgenic lines grown for 60 days in rhizotrons filled with vermiculite and nutritious soil (1:1), overview of RSA by EZ-RHIZO analysis of images, scale bar = 6cm (A). Total root number(B), number of links(C), average length(D), average diameter(E), average projected area(F), average surface area(G) of wild-type and transgenic lines grown normally for 2 months in rhizotrons. (Asterisks indicate a statistically significant difference compared with ZH11. \*\**P<0.01*, \*\*\**P <0.001*, \*\*\*\**P<0.0001*; One-way ANOVA).

### Exogenous auxin can restore the primary root length and lateral root number of *osaai1*

Numerous studies have shown that exogenously applied IAA can restore growth defects in plants[22–24]. To investigate whether external IAA affects the development of primary and lateral roots, different concentrations of IAA (0.1nM, 1nM, 10nM) were applied to *osaai1*, ZH11 and OE19 lines. The results showed that compared with the treatment without IAA, 0.1nM IAA treatment did not have significant effects on each line in plant height, and there were significant differences in primary root length, lateral root number and lateral root density between the wild type and transgenic lines, among which OE19 had the best growth. Under 1nM IAA treatment, there were no significant differences in plant height, primary root length, and lateral root density between the wild type and transgenic lines, but the number of lateral roots in *osaai1* was significantly less than that in ZH11 and OE19. Under 10nM IAA treatment, there were no significant differences in plant height, primary root length, lateral root number and lateral root density between the wild-type and transgenic lines. The above results indicated that the low concentration of IAA (1nM) treatment could compensate for the slow growth of the primary root of *osaai1*, but not for the differences in the growth of lateral roots among the lines; while the high concentration of IAA (10nM) treatment could compensate for the defects in the growth of the primary and lateral roots of *osaai1*(Fig3A-G, S3FigA-G and S4FigA, B). We stained the lateral root primordia of each line grown for 4 days with exogenously applied different concentrations of IAA (0, 0.1, 1, and 10nM), and the results were consistent with the phenotype described above, with no significant difference between the number of lateral root primordia of the wild-type and transgenic lines when exogenously applied 10nM IAA(S5FigA-E). This result suggests that exogenously applied IAA can restore the root growth defect caused by the *OsAAI1* mutation.

**Fig 3.**
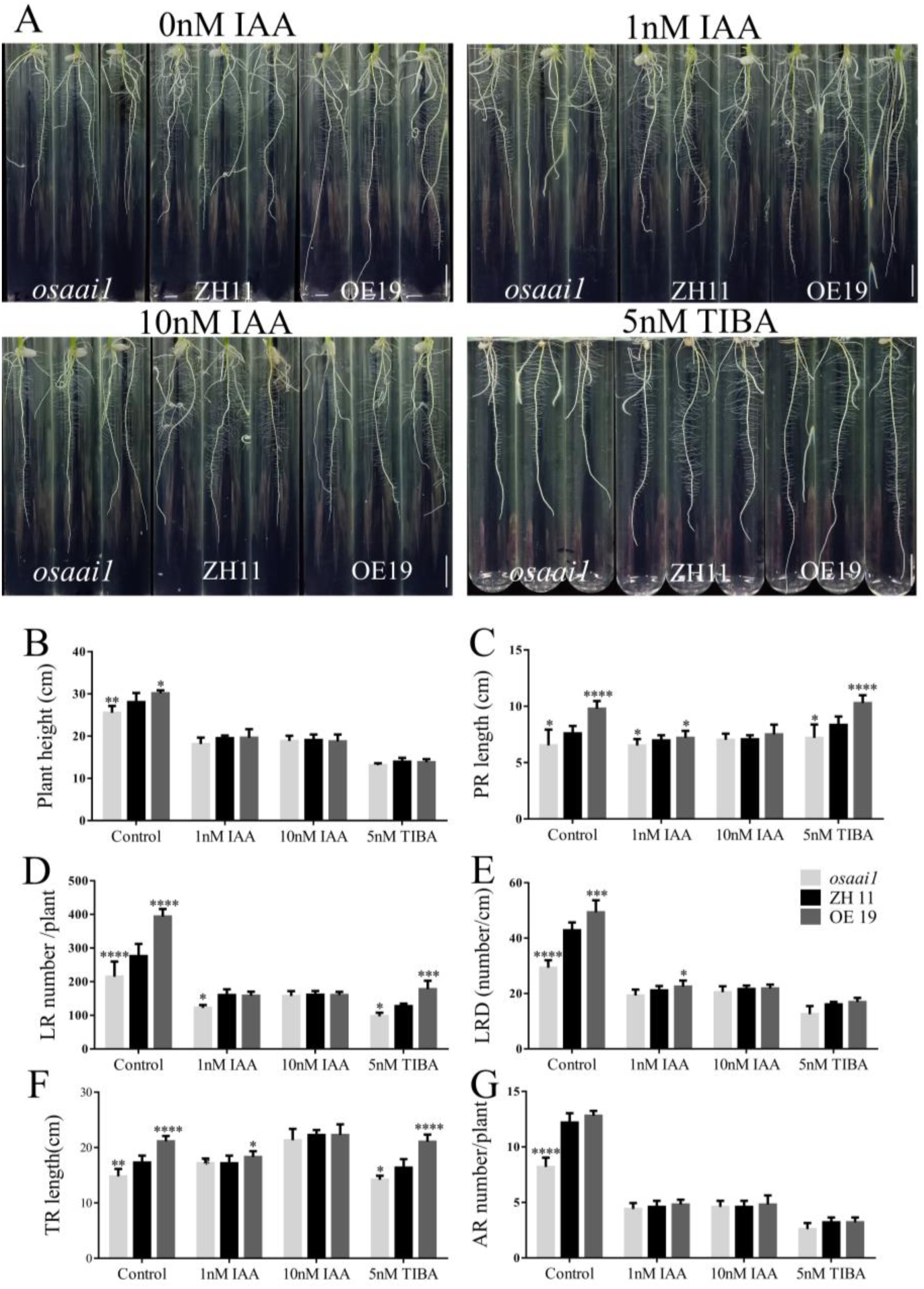
Phenotypic and statistical analyses of wild-type and transgenic lines treated with 0, 1, 10nM IAA and 5μM TIBA for 14 days. Phenology(A), scale bar = 1.5cm, plant height(B), primary root length(C), lateral root number(D), lateral root density(E), total root length(F) and adventitious root number(G) of wild type and transgenic lines treated with 0, 1, 10nM IAA and 5μM TIBA for 14 days. (Asterisks indicate a statistically significant difference compared with ZH11. **P<0.01, ***P <0.001, ****P<0.0001; One-way ANOVA).

As an inhibitor of auxin (IAA and IBA), 2,3,5-triiodo benzoic acid (TIBA) prevents polar auxin transport between cells and reduces auxin outflow[25]. It also competes directly with IAA at the efflux carrier and thus stops the transport of IAA in plants by occupying auxin transport channels[26]. By treating each line with exogenous TIBA (5nM, 5μM) for 14 days, no significant differences existed between the wild-type and transgenic lines in plant height and adventitious root number under the 5nM TIBA treatment compared with the treatment without TIBA, but significant differences existed between primary root length, lateral root number, lateral root density, and total root length, with *osaai1* growing the worst, and OE19 was the best (Fig3A-G). Under 5μM TIBA treatment, the wild-type and transgenic lines were severely inhibited, resulting in the inability of lateral roots to grow (S6FigA-G and S7Fig). We found that the 5nM TIBA treatment promoted the growth of primary root length and inhibited the growth of lateral root number, adventitious root number and adventitious root length of each line compared to the treatment without TIBA, 5μM TIBA treatment inhibited the growth of the entire root system of each line. Therefore, we conjecture that TIBA may have affected the transport of IAA from the lower part of the ground to the above ground, resulting in uneven distribution of IAA in the roots, which can only satisfy the normal growth of the primary roots in priority. However, its specific cause needs to be further investigated. Taken together, IAA was able to restore the root growth defect caused by the down-regulated expression of *osaai1*, suggesting that *OsAAI1* may affect rice root growth and development through the IAA signaling pathway.

### OsAAI1 involved in IAA signaling pathway

The ability of exogenous IAA to restore root development defects in *osaai1* suggests that *OsAAI1* may be involved in influencing changes in the IAA content of rice. Thus, we measured IAA content of the wild type and transgenic lines and showed that the IAA content of OE19 was significantly increased compared with ZH11, whereas the IAA content of *osaai1* was significantly decreased (Fig4A, B).

**Fig 4.**
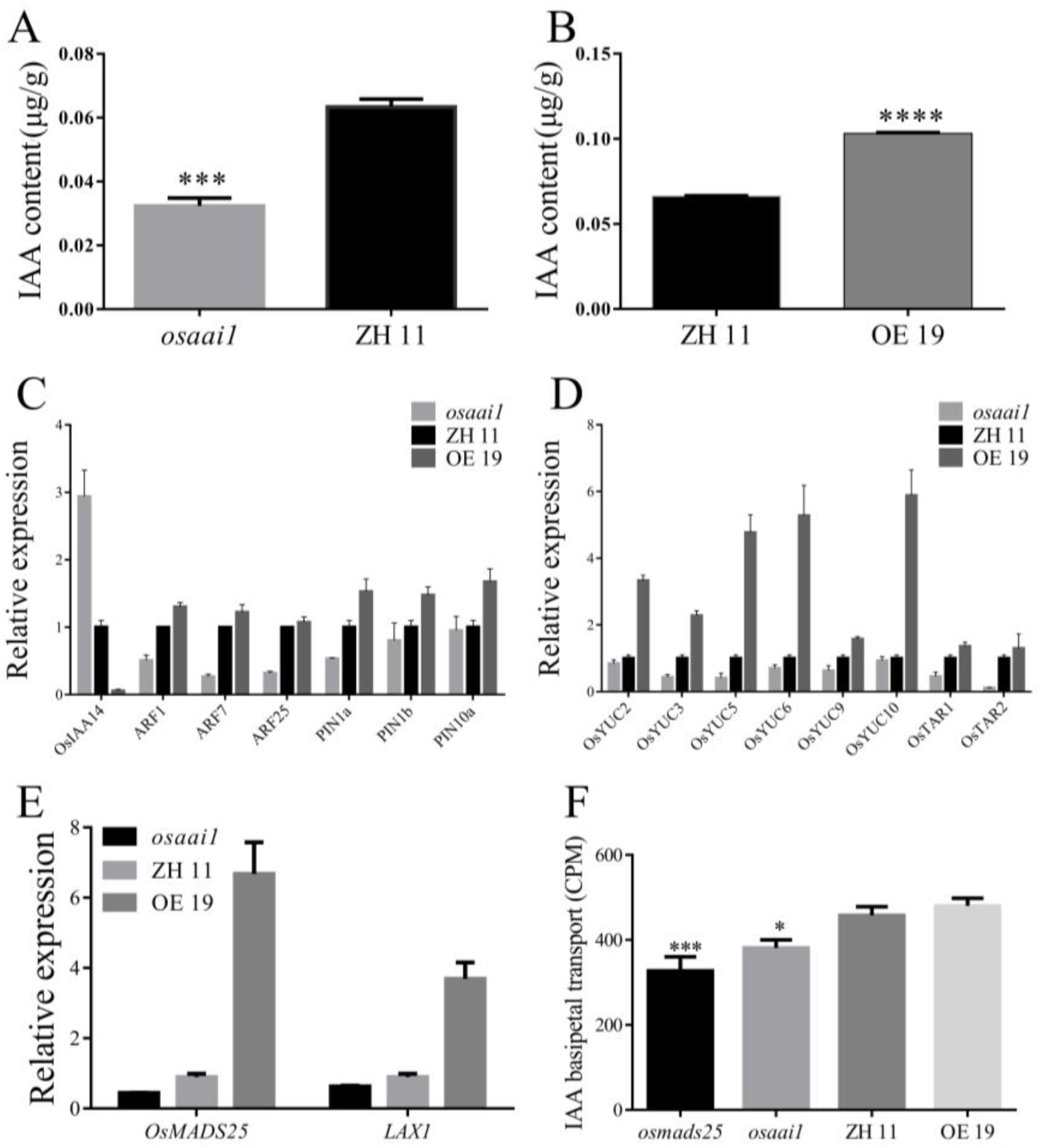
Analysis of IAA content, relative expression of IAA-related genes and IAA basipetal transport of wild-type and transgenic lines. IAA content of wild-type and transgenic lines (A, B). qRT-PCR of genes related to IAA biosynthesis, transport and metabolic pathways(C, D). Relative expression of *OsMADS25* and *LAX1* in wild-type and transgenic lines(E). IAA basipetal transport (CPM) of wild-type and transgenic lines(F). (Asterisks indicate a statistically significant difference compared with ZH11. \*\**P<0.01*, \*\*\**P <0.001*, \*\*\*\**P<0.0001*; One-way ANOVA.)

The accumulation of IAA in plants is closely related to IAA biosynthesis, transport, and degradation. To investigate whether *OsAAI1* regulates IAA content in rice by regulating auxin biosynthesis, perception and transport pathways, we examined the expression levels of auxin biosynthesis genes (*OsYUCs* and *OsTARs*), auxin transport genes (*OsPINs*) and auxin response factors genes (*ARFs*) by qRT-PCR. The results showed that the expression levels of IAA synthesis genes *OsYUC2*, *OsYUC3*, *OsYUC5*, *OsYUC6*, *OsYUC9*, *OsYUC10*, *OsTAR1* and *OsTAR2* were significantly higher in OE19 compared with ZH11, while they were significantly lower in *osaai1*(Fig4D); the expression levels of IAA response factors genes *ARF1*, *ARF7* and *ARF25* were increased in OE19, but decreased in osaai1(Fig4C); meanwhile, the expression levels of IAA transport genes *PIN1a*, *PIN1b*, and *PIN10b* were increased in OE19, but decreased in *osaai1*(Fig4C). We analyzed the expression levels of Aux/IAA family transcriptional repressors, the expression levels of *OsIAA14* were increased in *osaai1*, but decreased in OE19(Fig4C). To verify changes in auxin response, we conducted a comparison of polar auxin transport (PAT) activities in etiolated coleoptiles between transgenic lines and wild type. Our findings revealed a decrease in PAT activity to around 30% of wild type levels in osaai1 coleoptiles. Conversely, the PAT activity was slightly higher by 5% in OE19 when compared to the wild type (Fig4F). Taken as a whole, these findings indicate that *OsAAI1* controls the growth of rice roots through the regulation of IAA biosynthesis, perception and transport.

### Overexpression of *OsAAI1* improves root system development under drought stress

Our previous studies have shown that overexpression of *OsAAI1* can improve the drought tolerance of rice[21]. To investigate whether the enhancement of rice drought tolerance by overexpression of *OsAAI1* is related to root development, we observed the root growth of wild-type and transgenic lines treated with 3 months of water-off drought treatment. The results showed that OE19 had the widest root distribution and the highest total root number after two months of water cutting drought treatment, while the opposite was true for *osaai1*(Fig5, S8FigA-G and Table 2). This indicates that the more roots and wider roots distribution of OE19 allowed the plants to retain more water when subjected to drought stress, thus making them more resistant to drought stress.

**Fig 5.**
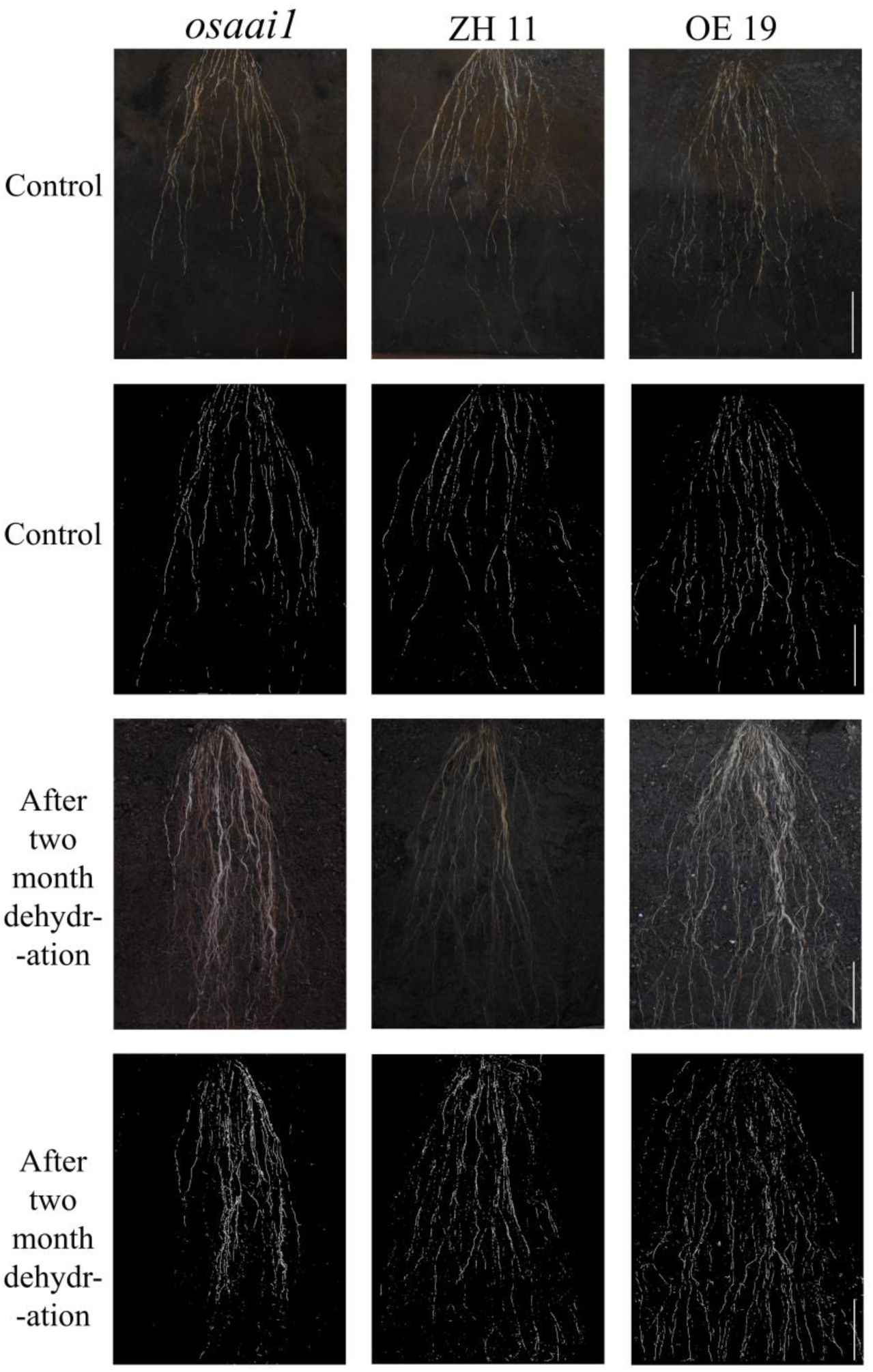
Root system architecture (RSA) of wild type and transgenic lines grown in rhizotrons with water-off drought treatment. Root system architecture (RSA) of wild type and transgenic lines grown for 60 days in rhizotrons filled with vermiculite and nutritious soil (1:1) supplemented with water-off drought treatment, an overview of RSA by EZ-RHIZO analysis of images, bar = 6cm.

**Table2.**
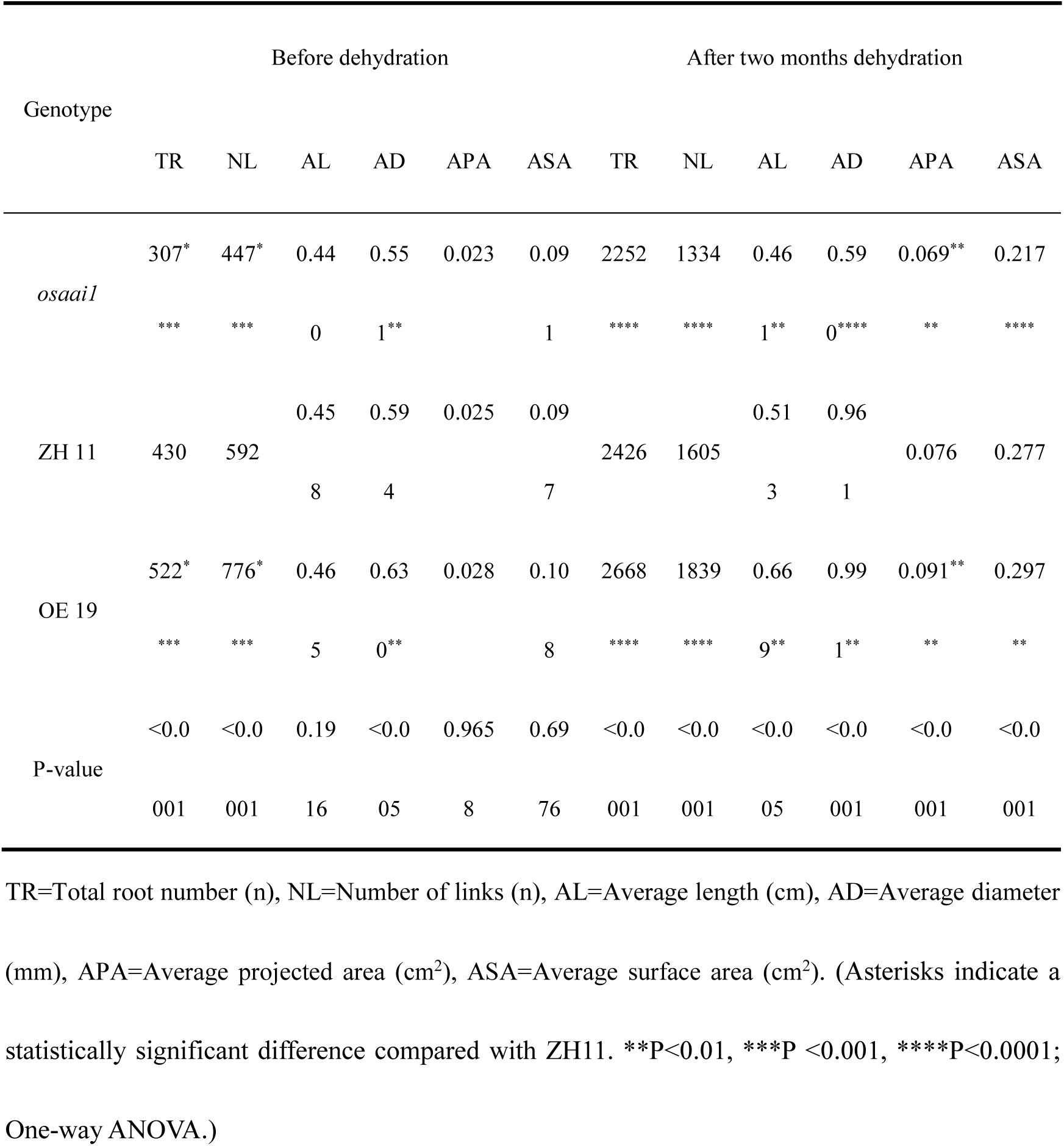
Pre-drought and two-month drought trait statistics for wild-type and transgenic lines.

### Exogenous application of IAA weakens the growth inhibition of *osaai1* by drought **stress**

Our previous study showed that *osaai1* growth was suppressed when 10% PEG was used to simulate drought treatment, and OE19 had the best growth status under 10% PEG treatment[21]. Our above results showed that the primary root length, lateral root number, lateral root density, total root length, and adventitious root number of OE19 were superior to those of ZH11 and *osaai1* under normal treatment, and there was no significant difference between the primary root length, number of lateral roots, lateral root density, and number of adventitious roots of the wild-type and transgenic lines when exogenous application of 10nM IAA was applied, suggesting that the exogenous application of IAA could compensate for the growth defects caused by *osaai1* mutation. We thus conjectured whether exogenous application of IAA could also attenuate the inhibition of *osaai1* by drought treatment. To verify this conjecture, we treated the wild-type and transgenic lines simultaneously with 10nM IAA and 10% PEG, and the results showed that the growth of all the lines was inhibited by 10% PEG and 10nM IAA treatment compared with 10% PEG treatment, but the growth of primary root length, lateral root number, lateral root density, and number of adventitious roots of the wild-type was not significantly different from those of the transgenic lines(Fig6A-G). We measured physiological indices of each line under different treatments. The results obtained were consistent with the above phenotypes (S9FigA-F). It indicates that exogenous application of IAA can attenuate the inhibition of *osaai1* by drought stress.

### OsAAI1 Physically Interacts with OsMADS25

We screened the yeast library and concluded that OsAAI1 interacts with OsMADS25, and we used the yeast two-hybrid system to test whether OsAAI1 interacts with OsMADS25 protein. We introduced the full length of OsAAI1 into the Gal4 activation domain of the prey vector (AD-OsAAI1), we fused the full-length of OsMADS25 to the Gal4 DNA binding domain of the bait vector (BD-OsMADS25). The bait and prey vectors were cotransformed into yeast and the protein-protein interaction was reconstructed (Fig7A). The OsAAI1–OsMADS25 interaction in plants was further verified by bimolecular fluorescence complementation (BiFC) assays. For the BiFC assays, OsAAI1 was fused to a C-terminal yellow fluorescent protein (YFP) fragment (OsAAI1-cYFP), and the OsMADS25 protein was fused to an N-terminal YFP fragment (OsMADS25-nYFP). When fused OsAAI1-cYFP was coexpressed with OsMADS25-nYFP in leaves of *Nicotiana benthamiana*, a YFP fluorescence signal-as revealed by staining with 4′,6-diamidino-2-phenylindole-was observed in transformed cell nuclei (Fig7C). No fluorescence was detected in the negative control experiments (Fig7C). Further verified with firefly luciferase complementation imaging (LCI) assay (Fig7B), All results demonstrate that the OsAAI1 physically interacts with the OsMADS25 transcription factor.

**Fig 6.**
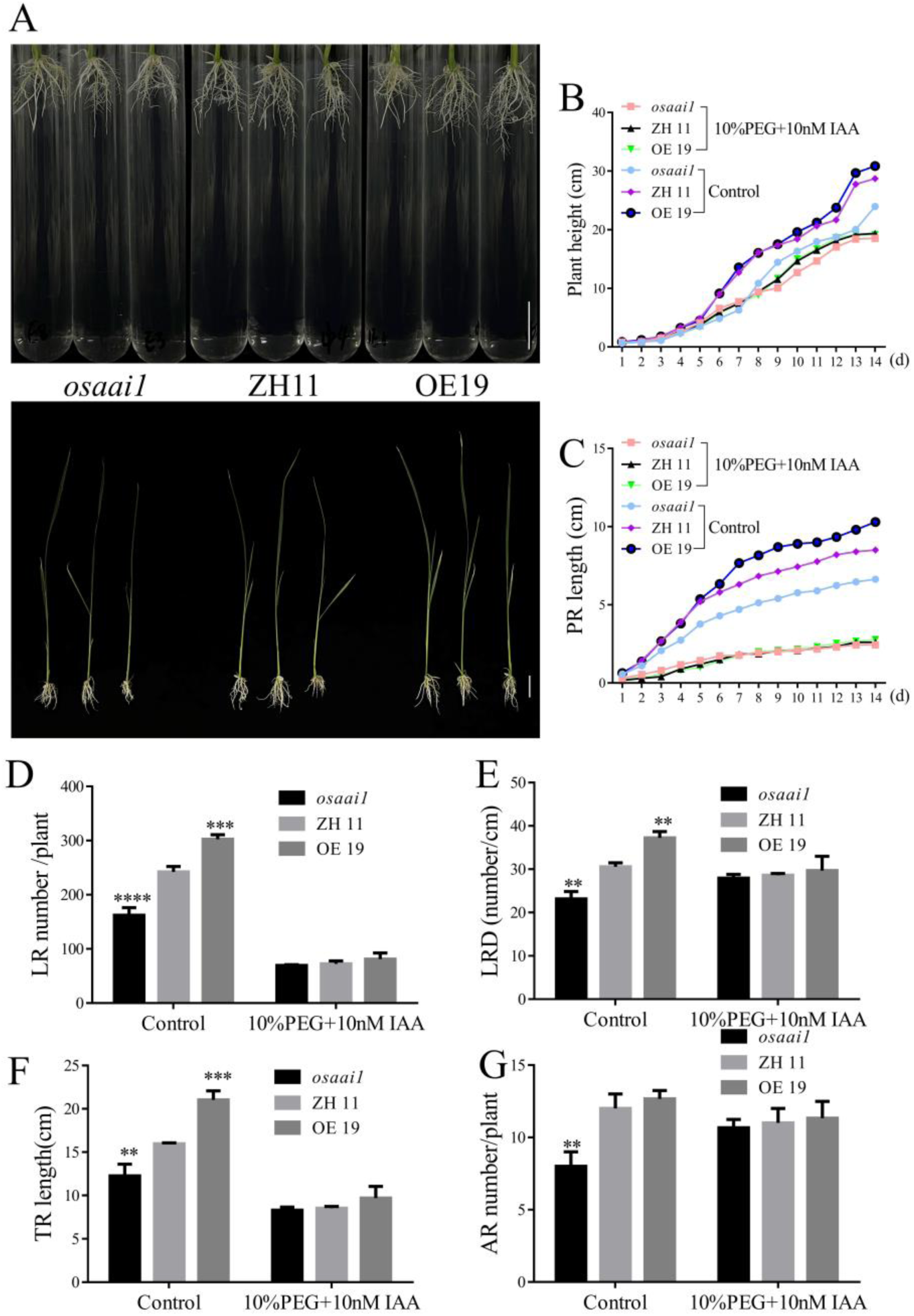
Phenotypic and statistical analyses of wild-type and transgenic lines treated with 10%PEG + 10nM IAA. Phenology(A), bar = 1.5cm, plant height(B), primary root length(C), lateral root number(D), lateral root density(E), total root length(F) and adventitious root number(G) of wild type and transgenic lines treated with 10%PEG + 10nM IAA for 14 days. (Asterisks indicate a statistically significant difference compared with ZH11. \*\**P<0.01*, \*\*\**P <0.001*, \*\*\*\**P<0.0001*; One-way ANOVA).

**Fig 7.**
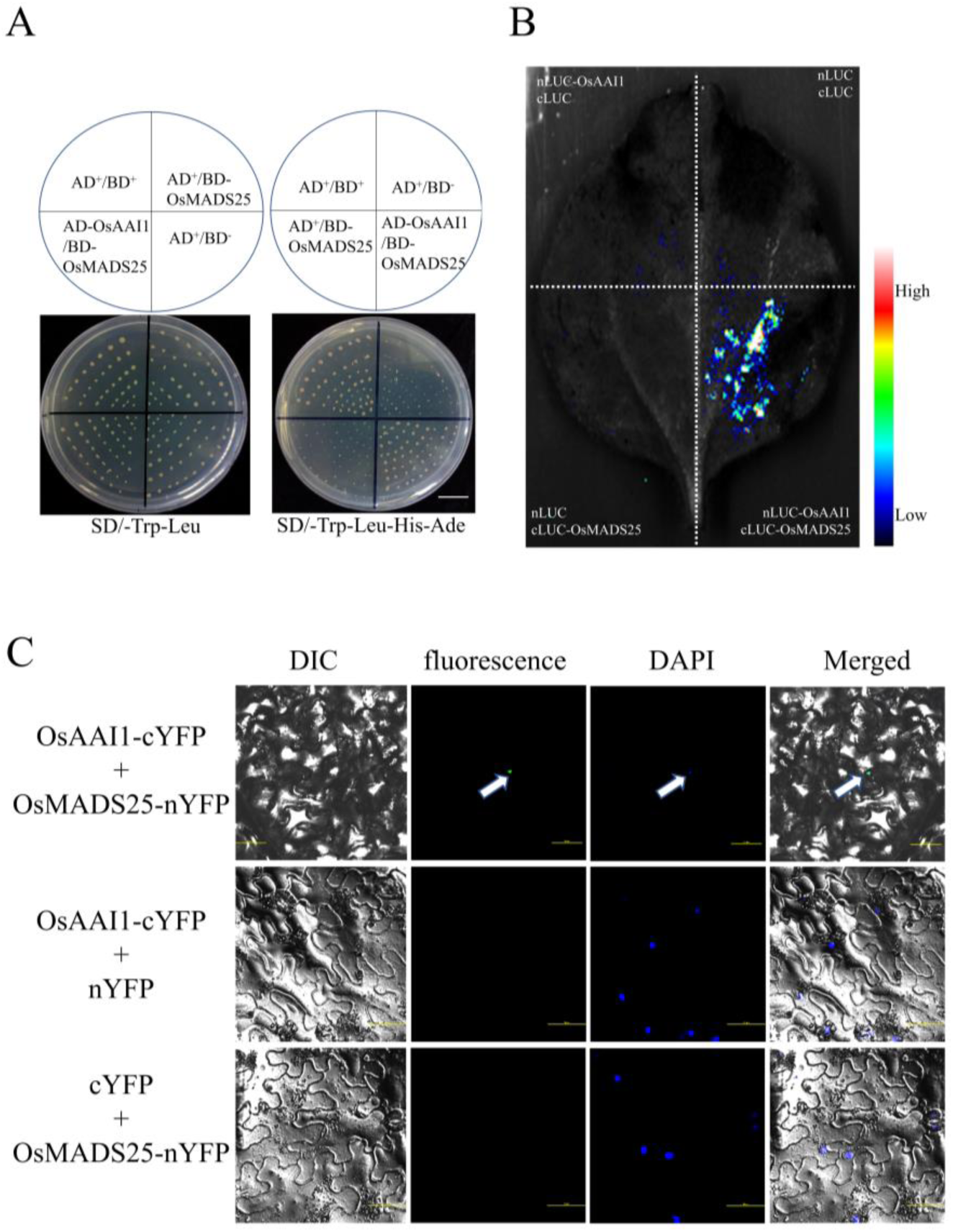
OsAAI1 Physically Interacts with OsMADS25. Y2H assays showing the interaction between OsAAI1 and OsMADS25, the empty pGADT7 and pGBKT7 vector was used as a negative control, scale bar = 1.5cm(A). Firefly luciferase complementation imaging (LCI) assay of the interaction of OsAAI1 with OsMADS25 (B). OsAAI1 and OsMADS25 bimolecular fluorescence complementary experiment analysis, scale bar = 50μm(C).

### Exogenous application of IAA weakens the growth inhibition of *osmads25* by drought stress

The current studies on the stress tolerance of *OsMADS25* gene mainly focus on salt tolerance and cold tolerance[27–29]. While the drought resistance aspect has not been reported so far. To verify the drought resistance of *OsMADS25*, we treated *osmads25* with 10% PEG for 14 days, observed the phenotype and performed statistical analysis of the phenotypic data. The results showed that the growth of both *osmads25* and ZH11 was significantly inhibited under 10% PEG treatment, but the plant height, primary root length, number of lateral roots, lateral root density, and number of adventitious roots of ZH11 were significantly better than those of *osmads25*(Fig8A-G), suggesting that the *osmads25* was more sensitive to drought stress.

**Fig 8.**
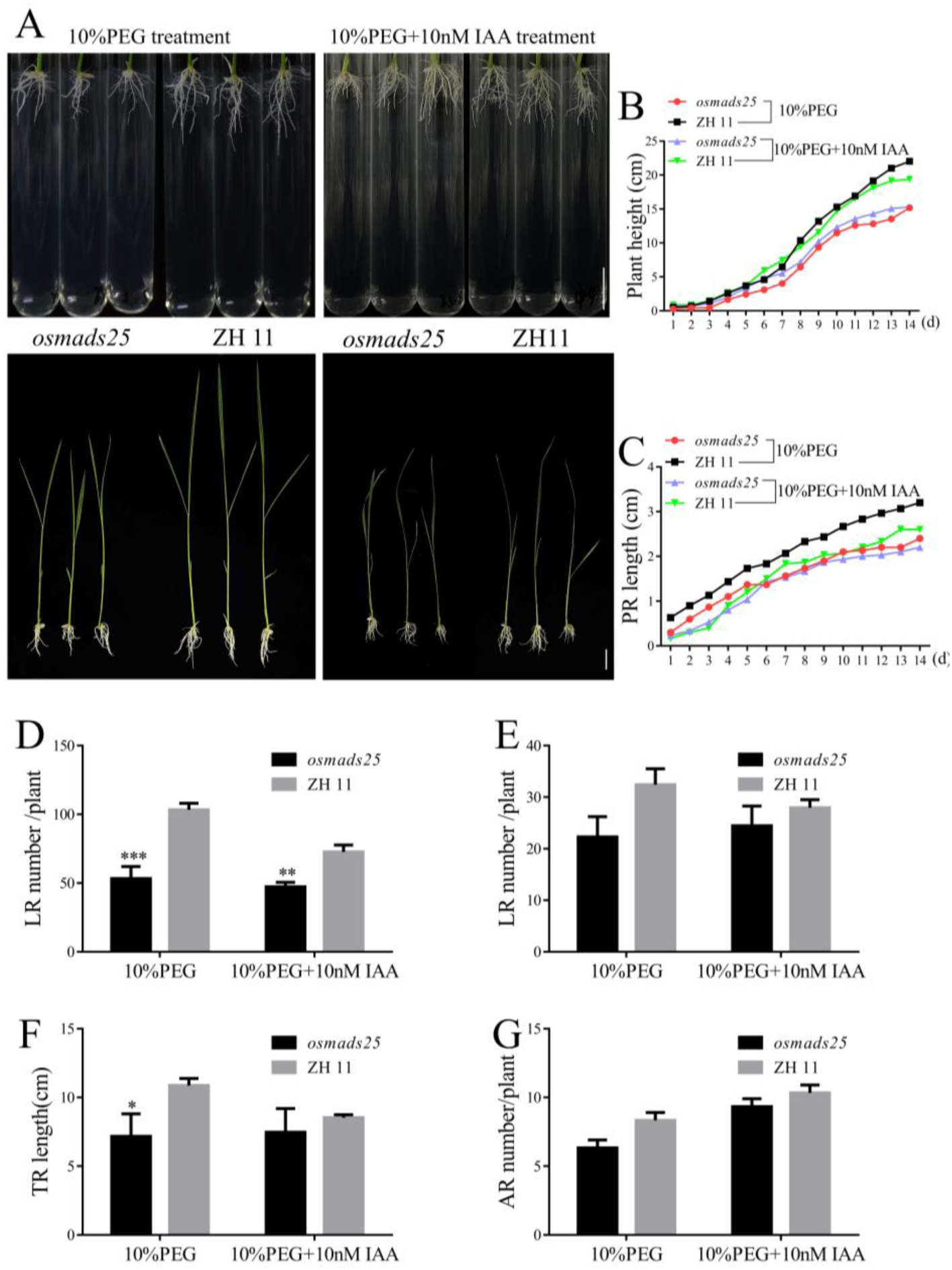
Phenotypic and statistical analyses of wild-type and transgenic lines treated with 10%PEG, 10%PEG + 10nM IAA. Phenology(A), scale bar = 2cm, plant height(B), primary root length(C), lateral root number(D), lateral root density(E), total root length(F) and adventitious root number(G) of wild type and *osmads25* treated with 10%PEG, 10%PEG + 10nM IAA for 14 days. (Asterisks indicate a statistically significant difference compared with ZH11. \*\**P<0.01*, \*\*\**P <0.001*, \*\*\*\**P<0.0001*; One-way ANOVA)

Previous studies showed that overexpression of *OsMADS25* significantly enhanced primary root length and lateral root density in rice, and exogenously applied auxin was able to restore the root growth and developmental defects of OsMADS25-RNAi[30]. To verify whether exogenous IAA could also alleviate the suppression of *osmads25* by drought stress, we treated *osmads25* and ZH11 simultaneously with 10nM IAA and 10% PEG, and the results showed that the growth of all lines was suppressed under 10% PEG and 10nM IAA treatment compared with 10% PEG treatment, but exogenous application of IAA narrowed the differences between *osmads25* and ZH11. We measured physiological indices of each strain under different treatments. The results obtained were consistent with the phenotypes described above (S9FigG-L). It indicates that exogenous application of IAA can also attenuate the inhibition of *osmads25* strain by drought stress.

### OsAAI1 with OsMADS25 to regulate the expression of downstream genes

Previous studies have shown that *OsMADS25* binds to the CArG-box in the promoter region of *OsIAA14* and regulates LR formation[30]. *OsMADS25* enhances growth hormone biosynthesis through activation of *OsYUC4*, which promotes growth hormone signaling to regulate root development[27]. To further explore the mechanism of interaction between OsAAI1 and OsMADS25, we screened four downstream target genes of *OsMADS25* (*OsIAA14*, *OsYUC4*, *LAX1*, *OsBAG4*). The full-length cDNA of *OsMADS25* was fused to the GAL4 activation structural domain of the pGADT7 vector, and the promoter sequences of *OsBAG4* and *LAX1*, which carry CArG-box motifs, were ligated into the pHISi vector to performed yeast one hybrid (Y1H) assay. The result showed that *OsMADS25* directly interacted with the *OsBAG4* and *LAX1* promoter sequences (Fig9A, B). Then, EMSA analysis was conducted to investigate the DNA binding specificity of OsMADS25 towards the *OsBAG4* and *LAX1* promoters containing CArG-box motifs. The results in Fig 9C demonstrate a distinct band shift when probe from the *OsBAG4* and *LAX1* promoter region was exposed to OsMADS25-His protein. Subsequent competition assays further validated the specificity of these binding interaction (Fig9C). To further characterize the roles of *OsMADS25* and *OsAAI1* in regulating the expression of *OsIAA14*, *OsYUC4*, *LAX1*, and *OsBAG4*, we performed a transient expression assay of the reporter gene LUC in *Nicotiana benthamiana* leaves. The effectors were *OsMADS25* and *OsAAI1* driven by the CaMV35S promoter. The promotor (pro) sequences of *OsIAA14*, *OsYUC4*, *LAX1*, and *OsBAG4* containing CArG-box cis-elements were fused to the LUC gene and used as reporter genes. The plasmids carrying the reporter and effector genes were co-infiltrated into the leaves of *N. benthamiana*, and the LUC activity assay showed that *OsMADS25* and *OsAAI1* repressed the expression of the LUC reporter genes driven by the *OsIAA14* promoter, and promoted the expression of the LUC reporter genes driven by the promoters of *OsYUC4*, *LAX1*, and *OsBAG4* (Fig9D).

### Phenotypic trait analysis of *osaai1/osmads25*

To further investigate the mechanism of interaction between OsAAI1 and OsMADS25, we obtained hybrid seeds(*osaai1/osmads25*) of *osaai1* mutant line and *osmads25* mutant line by means of artificial hybridization and performed phenotypic data on *osaai1/osmads25*, *osaai1*, *osmads25*, ZH11, and OE19 lines with 14 days of normal growth statistics and analysis. The results showed that OE19 had the optimal plant height, primary root length, number of lateral roots, lateral root density, total root length, and number of the adventitious roots compared to ZH11, while *osaai1/osmads25* was the poorest growing of all the lines, followed by *osmads25* and *osaai1*(Fig10A-G), indicating that the *osaai1/osmads25* double mutation severely affected the normal growth of the plants.

## Discussion

### *OsAAI1* regulates primary root elongation by modulating the length of rice primary root cells

*OsAAI1* is a member of the HPS-like subfamily of the AAI_LTSS superfamily, and members of this subfamily play important roles in helping plants resist abiotic stresses. Our previous study showed that *OsAAI1* enhances drought tolerance in rice through the ABA-regulated pathway and the ROS-scavenging pathway[21]. However, functional studies of *OsAAI1* in rice root development have not been reported. In this study, we found that overexpression of *OsAAI1* increased rice primary root length, lateral root number, lateral root density, adventitious root number, and total root length compared with the wild type, whereas the opposite was true for the mutant line (Fig1A-F). Lateral root primordia staining showed that the number and density of lateral root primordia on the primary roots of *osaai1* were significantly lower than those of ZH11, whereas those of OE19 were significantly higher than those of ZH11(Fig1G, H). Transverse sectioning of paraffin sections showed that OE19 had more thin-walled cells in the xylem and phloem and were more tightly arranged than ZH11 and *osaai1*. Moreover, the lateral root primordia appeared first in OE19(S1FigA). Longitudinal sectioning of paraffin sections showed that the cells in the elongation zone of OE19 were longer compared with those of ZH11 and *osaai1*(Fig1I and S1FigB). Rhizotron experiment showed that, among the wild type and transgenic lines that had been grown normally for 1-3 months, OE19 consistently had the best root growth, followed by the ZH11, and the worst one of *osaai1*(Fig2A-G and S2Fig). In summary, *OsAAI1* regulates the elongation of primary roots and the formation of lateral root primordia by modulating the length of rice primary root cells.

### *OsAAI1* regulates root system development via auxin signaling pathway

The plant hormone IAA plays a crucial role in the plant’s growth and development [22–24]. In this study, we showed that *OsAAI1* appears to promote root development by regulating auxin levels. Firstly, The IAA content of OE19 was significantly higher than that of ZH11 and *osaai1*(Fig4A, B). Secondly, exogenous application of IAA could compensate for the root growth defects induced by *OsAAI1* mutation (Fig3A-G, S3FigA-G and S4FigA, B), suggesting that *OsAAI1* may affect rice root growth and development through IAA signaling pathway. There are two main pathways for growth hormone synthesis: tryptophan-dependent pathway and non-tryptophan-dependent pathway. The tryptophan-dependent pathway is divided into three main branches according to the intermediates of growth hormone IAA synthesis: the indolee-3-pyruvate acid (IPA) pathway, the tryptamine (TAM) pathway, and the indolee-3-acetonitrile (IAN) pathway in *Arabidopsis*[31]. In the IPyA pathway, tryptophan aminotransferases (TAA1/TARs) synthesize IPyA from L-tryptophan[32,33], and YUCCAs (YUCs) convert IPyA to IAA[31,34]. Kakei et al. demonstrated that *OsTAR1* catalyzes the conversion of L-tryptophan to IPyA[35]. *OsTAR2* may play a major role in ethylene response of rice root[36]. The YUC family of flavin monooxygenases are the enzymes for the final step of the IPA biosynthesis pathway in Arabidopsis[31,37]. *OsYUC2* was involved in IAA synthesis, overexpression of *OsYUC2* in roots influenced root growth and development[38]. Overexpression of *the YUC3* gene in shoots led to an auxin overproduction phenotype in shoots[39]. *YUC6* was suggested to be mainly responsible for auxin biosynthesis in shoots[33]. *YUC9* and *YUC* gene family members redundantly mediate cut-induced auxin biosynthesis in Arabidopsis[40]. In our study, the expression levels of IAA synthesis genes *OsYUC2*, *OsYUC3*, *OsYUC5*, *OsYUC6*, *OsYUC9*, *OsYUC10*, *OsTAR1* and *OsTAR2* were significantly higher in OE19 compared with ZH11, while they were significantly lower in *osaai1*(Fig4D). Auxin response factors (*ARFs*) bind auxin response promoter elements and mediate transcriptional responses to auxin[37]. *ARF1* and *ARF2* regulate senescence and floral organ abscission in *Arabidopsis thaliana*[41]. *ARF7* and *ARF19* promote leaf expansion and auxin-induced lateral root formation[42]. In our study, the expression levels of IAA response factors genes *ARF1*, *ARF7* and *ARF25* were increased in OE19, but decreased in *osaai1*(Fig4C). The auxin efflux carrier PIN-FORMED (*PIN*) family is one of the major protein families involved in Polar auxin transport (PAT)[43]. LPA1 might control auxin transport via the initiation of PIN1a to increase planting density and activate plant defense gene expressions[44].*OsPIN1b* was mainly expressed in roots, stems and sheaths at the seedling stage, Mutation of *OsPIN1b* resulted in IAA homeostasis disturbed[45]. In our study, the expression levels of IAA transport genes *PIN1a*, *PIN1b*, and *PIN10b* were increased in OE19, but decreased in *osaai1*(Fig4C). The expression levels of *OsIAA14* were increased in *osaai1*, but decreased in OE19(Fig4C). In summary, these results suggest that *OsAAI1* regulates rice root development by regulating IAA biosynthesis, response and transport.

### OsAAI1 interacts with OsMADS25 to regulate the expression of downstream genes and ultimately affects ROS homeostasis and IAA signaling pathways

*OsMADS25* is a member of the AGL17 clade of homologs in rice[46]. Previous studies have shown that *OsMADS25* binds to the CArG-box in the promoter region of *OsIAA14* and regulates LR formation[30]. *OsMADS25* enhances growth hormone biosynthesis through activation of *OsYUC4*, which promotes growth hormone signaling to regulate root development[27]. We screened yeast libraries and found that there was a mutual interaction between OsAAI1 and OsMADS25 and demonstrated that by yeast two hybrid (Y2H), BiFC and split-luciferase complementation (SLC) assays(Fig7A-C). Yeast one-hybrid and EMSA assays showed that *OsMADS25* can directly interact with *OsBAG4* and *LAX1* promoter sequences (Fig9A-C). Transient expression assays of the reporter gene LUC showed that *OsMADS25* and *OsAAI1* repressed the expression of the LUC reporter gene driven by the *OsIAA14* promoter and promoted the expression of the LUC reporter gene driven by the *OsYUC4*, *LAX1*, and *OsBAG4* promoters (Fig9 D). This is consistent with previous findings[27,30], suggesting that OsAAI1 and OsMADS25 regulate the expression of downstream genes through interactions.

More importantly, the transcript levels of *OsMADS25* were significantly enhanced in OE19 and significantly reduced in *osaai1*(Fig4E), implying that *OsMADS25* may enhance the expression of *OsAAI1*. In previous study, *OsMADS25* directly activated the expression of the gene encoding the glutathione sulfotransferase *OsGST4*, an in vitro recombinant protein of *OsGST4* that has the ability to scavenge ROS, thus *OsMADS25* increased ROS scavenging and thus enhanced rice salt tolerance by directly activating the expression of *OsGST4*[27], whereas in our previous study, OE19 accumulated fewer ROS than ZH11 and *osaai1* under drought stress, suggesting that the ROS scavenging capacity of OE19 was enhanced[21]. In this study, we found that exogenous application of IAA restored the root developmental defects caused by *OsAAI1* mutation (Fig3A-G), and that exogenous application of IAA alleviated the suppression of the *OsAAI1* mutant line under drought stress. There were no significant differences between the growth phenotypes and physiological indices of the wild-type and transgenic lines when exogenously applying 10nM IAA under 10% PEG-simulated drought stress conditions (Fig 6A-G). There was also no significant difference between the activities of ROS-related scavenging enzymes (CAT, APX, GPX) (S9FigA-F). Therefore, we suggest that OsAAI1 may interact with OsMADS25 to regulate the expression of downstream target genes and enhance the ROS scavenging ability in response to the IAA signaling pathway, thereby enhancing the tolerance of rice to drought stress.

*LAX1* encodes a plant-specific bHLH transcription factor with pleiotropic effects. Apart from its role in controlling axillary bud primordia formation in rice, LAX1 also modulates auxin transport by physically interacting with OsPID, influencing rice flower organ development. Notably, *LAX1* plays a positive role in drought stress response and is a key candidate gene for drought resistance[47–50]. BAG (Bcl-2-associated athanogene) proteins, evolutionarily conserved in animals and plants, exhibit chaperone function. OsBAG4, a member of group I within this protein family, possesses a conserved ubiquitin-like structure. Overexpression of *OsBAG4* leads to stunted growth and heightened immune responses, enhancing disease resistance[51,52]. Similarly, overexpression of *AtBAG4* in tobacco boosts tolerance to various abiotic stresses[53]. Through yeast one-hybrid and EMSA assays, we demonstrated that OsMADS25 can bind to the conserved CArG box sequence in the promoter regions of *LAX1* and *OsBAG4*(Fig9A-C). Additionally, LUC activity assays revealed that the interaction between OsMADS25 and OsAAI1 enhances the transcriptional expression of *LAX1* and *OsBAG4*(Fig9D).

In summary, we summarized the mode of interaction between OsAAI1 and OsMADS25. When OsAAI1 was present, OsAAI1 forms protein complexes with OsMADS25. On the one hand, the protein complex inhibited the expression of *OsIAA14*, thereby affecting the signaling pathway of IAA. the protein complex promoted the expression of *OsYUC4*, which resulted in the elevation of endogenous IAA content in *OsAAI1* overexpression line, and the signaling pathway of IAA and the content of IAA jointly regulated rice root development, which resulted in better root development of *OsAAI1* overexpression line than that of wild-type and mutant line, thus enhancing the tolerance of rice to drought stress. On the other hand, OsAAI1 interacted with OsMADS25 to enhance the expression of *LAX1* and *OsBAG4*, which enhanced the tolerance of rice to drought stress (Fig 11).

**Fig 9.**
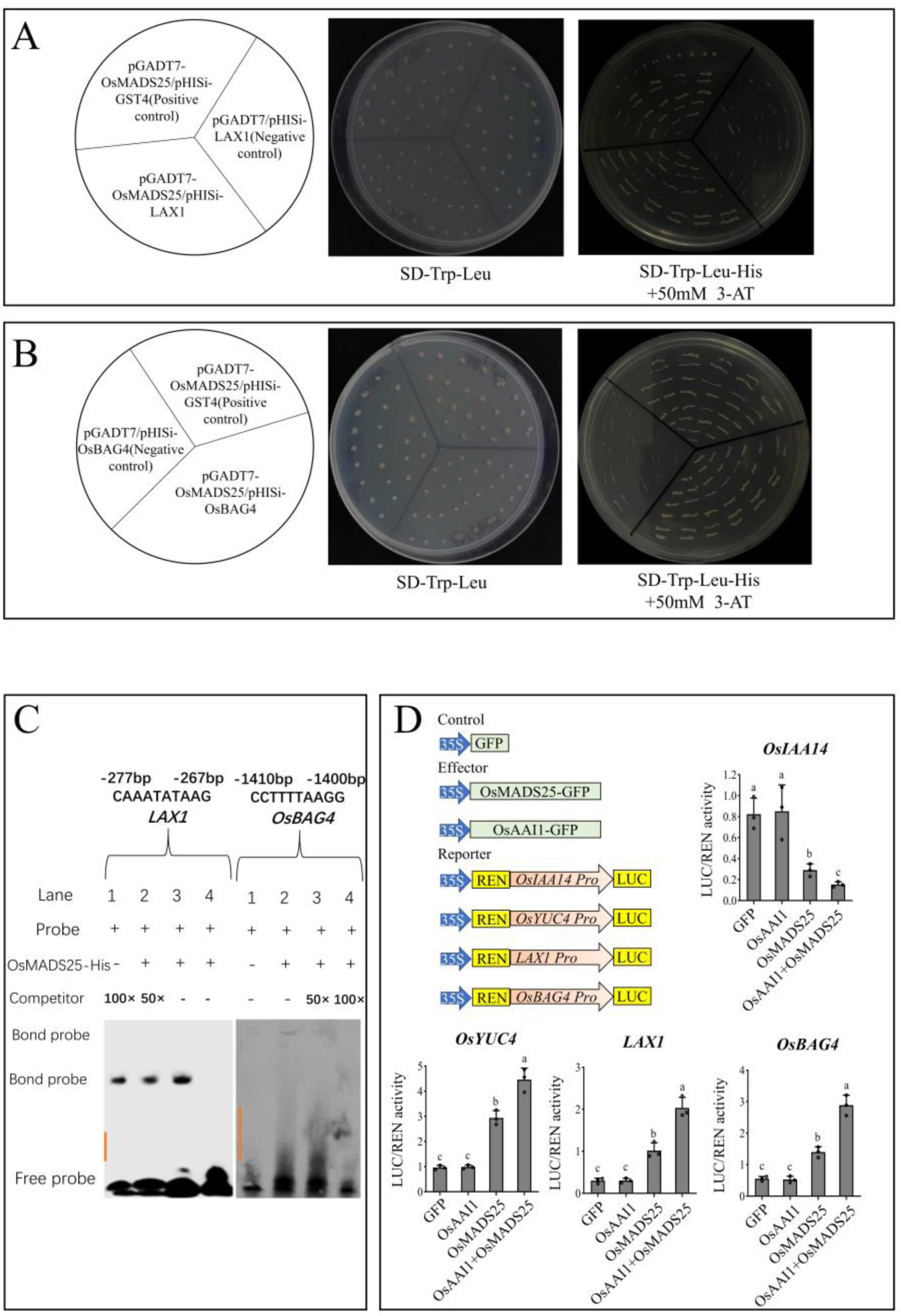
OsAAI1 with OsMADS25 to regulate the expression of downstream genes. The Yeast One Hybrid (Y1H) assay results of *OsMADS25* and *LAX1*, pGADT7-OsMADS25/pHISi-GST4 was used as positive controls and pGADT7/pHISi-LAX1 as negative control, scale bar=2cm (A). The Yeast One Hybrid (Y1H) assay results of *OsMADS25* and *OsBAG4*, pGADT7-OsMADS25/pHISi-GST4 was used as positive controls and pGADT7/pHISi- OsBAG4 as negative control. scale bar=2cm(B). Electrophoretic mobility shift assays (EMSA) indicating OsMADS25 binding specific CArG–box motifs(C). Schematic diagram of the effector and reporter used for transient trans-regulation studies.35S::OsMADS25 and 35S::OsAAI1 were used as effector genes OsIAA14pro::GUS, OsYUC4pro::GUS, LAX1pro::GUS, and OsBAG4pro::GUS were used as reporter genes, and 35S::GFP was used as a negative control. Statistical analysis of relative LUC activity (D).

**Fig 10.**
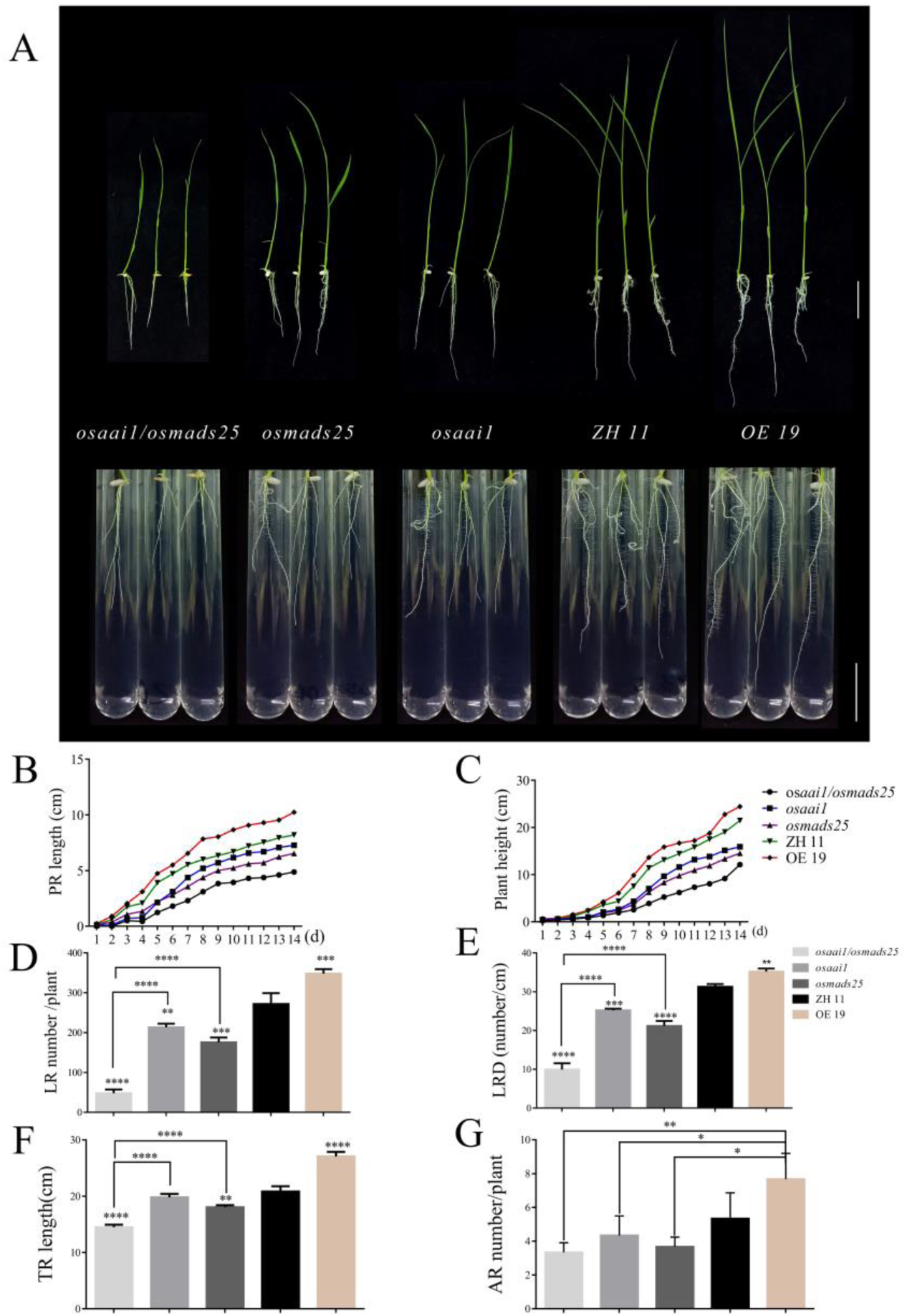
Phenotypic trait analysis of osaai1/osmads25. Phenotypes of *osaai1/osmads25*, *osaai1*, *osmads25*, ZH11 and OE19 grown normally for 14 days, scale bar = 2.5cm (a), primary root length(b), plant height(c), lateral root number(d), lateral root density(e), total root length(f) and adventitious root number(g) of wild type and transgenic lines at 14 days of normal growth. (Asterisks indicate a statistically significant difference compared with ZH11. \*\**P<0.01*, \*\*\**P <0.001*, \*\*\*\**P<0.0001*; One-way ANOVA.)

**Fig 11.**
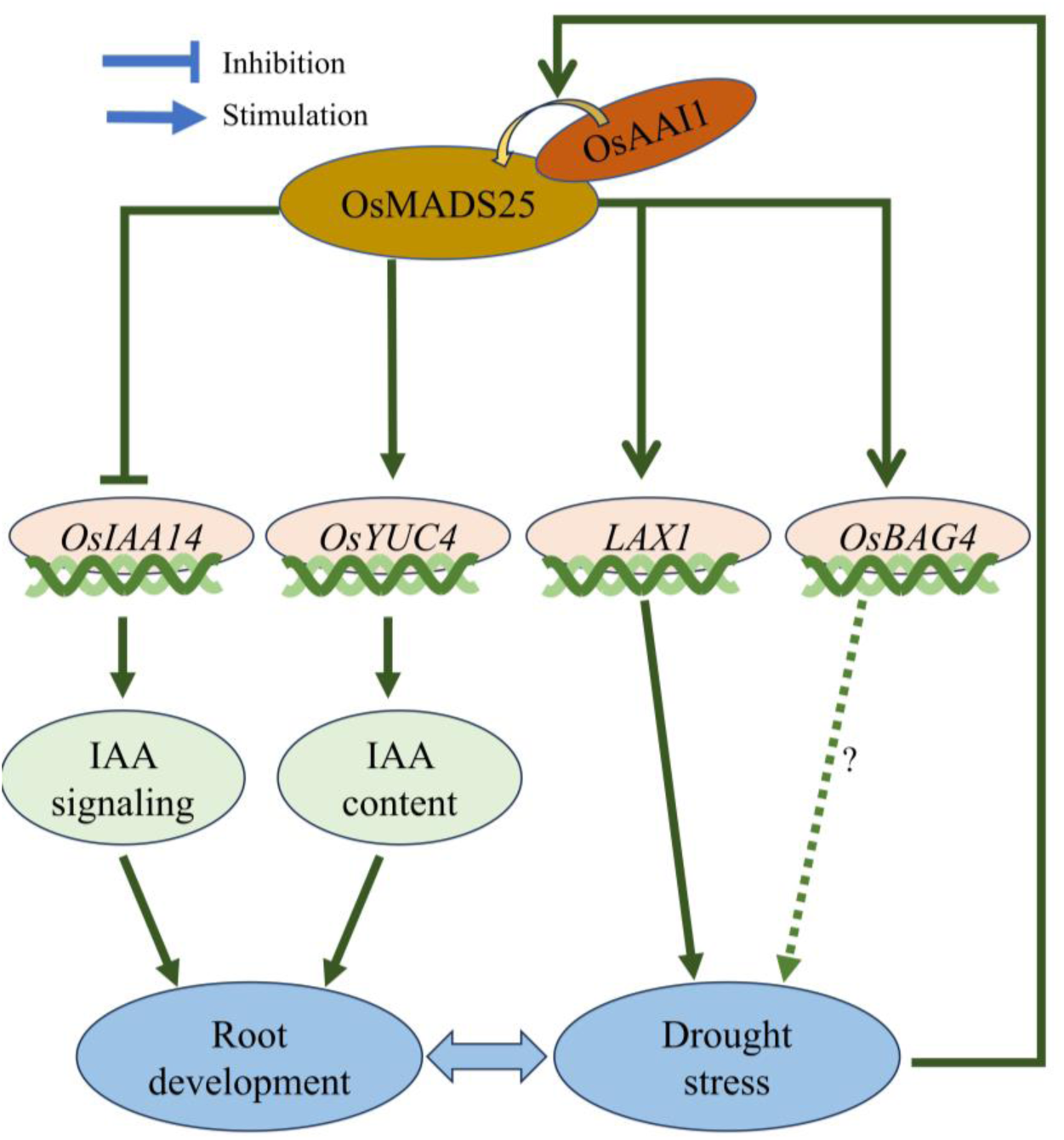
Mechanism of interactions between OsAAI1 and OsMADS25. The protein complex suppressed OsIAA14 expression, impacting the IAA signaling pathway; it also enhanced OsYUC4 expression, resulting in elevated endogenous IAA levels in the OsAAI1 overexpression strain. The coordinated modulation of the IAA signaling pathway and IAA levels influenced rice root growth, leading to enhanced root development in the OsAAI1 overexpression strain compared to the wild-type and mutant strains. This improved root growth contributed to increased drought resistance in rice. Furthermore, OsAAI1 interacted with OsMADS25 to stimulate *LAX1* and *OsBAG4* expression, further bolstering rice’s drought tolerance.

## Materials and methods

### Plant materials and hormone treatment

In this study, The Oryza sativa *L.* variety “Zhong Hua 11” was used as wild type for physiological experiments and genetic transformation. *OsAAI1* mutant line(*osaai1-1*) and overexpression line (OE19) were kept in this laboratory. Sensitivity analysis of IAA and TIBA of wild type and transgenic lines: IAA (0.1nM, 1nM, 10nM) and TIBA (5μM, 5nM) were added to the 1/2 MS medium at about 60°C. Then, the phenotypes of the three lines after hormone treatment were observed and statistically analyzed. Sensitivity analysis of 10nM IAA and 10%PEG of wild type and transgenic lines: 10nM IAA 10%PEG were added to the 1/2 MS medium at about 60°C. Then, the phenotypes of the three lines after hormone treatment were observed and statistically analyzed.

### Hybrid rice assay

Cut 3/5 of the glumes of rice before the opening of the glumes, spray warm water on them, the filaments absorb water and push the anthers out, pop the anthers, complete the operation of de-sexing, de-sexed rice as the female parent, bagging and fixing of the treated female parent, fixed with paper clips. Select the flowering rice as the parent for pollination, and after pollination, seal the bag and cultivate until maturity.

### Quantification of endogenous IAA

For IAA quantification, the plants of two-weeks-old ZH11, OE19 and *osaai1* were harvested and used for measurement of IAA by a liquid chromatography system (ACCHROM S3000) high performance liquid chromatograph, Alphasil VC-C18 (4.6mm * 250mm, 5μm) chromatography measure the contents of IAA. Each sample was repeated three times.

### Total RNA extraction and quantitative PCR analysis

Total RNA was isolated using Life’s Trizol reagent (code: 15596-026). The RNA extracted was then utilized for cDNA synthesis with Vazyme’s HiScript II qRT SuperMix for qPCR (+gDNA wiper) reagent Kit (code: R223-01). Subsequently, qPCR analysis was conducted on a Bio-Rad CFX96 instrument using Vazyme’s ChamQ Universal SYBR qPCR Master Mix reagent (code: Q711-02) as per the manufacturer’s guidelines. The housekeeping gene β-Actin from rice was employed as an internal control. Each experimental set included three independent biological replicates and three technical replicates. Details of the qRT-PCR primers can be found in S1Table.

### Yeast two-hybrid assays (Y2H)

For yeast two hybrid (Y2H) analysis, the coding sequence (CDS) of *OsAAI1* was amplified by PCR and inserted into the pGADT7 vector, and the coding sequence (CDS) of *OsMADS25* was amplified by PCR and inserted into the pGBKT7 vector. To examine whether there is a direct physical interaction between *OsAAI1* and *OsMADS25* in yeast (AH109). The primers used for the Y2H assay are listed in S2Table.

### Bimolecular fluorescence complementation assay

The *OsAAI1* full-length CDS sequence was cloned into the BiFC vector pAB855-cYFP vector, and the full-length CDS sequence of *OsMADS25* was cloned into the BiFC vector pAB862-nYFP vector. The constructed BiFC vectors OsAAI1-cYFP and OsMADS25-nYFP were transferred into the Agrobacterium GV3101 strain. The constructs were transiently coexpressed in tobacco leaves (*Nicotiana benthamiana*) performed as described previously[54]. The images were captured using a confocal laser-scanning microscope (Leica TCSSP8, Germany). The primers used for the BiFC assay are listed in S2Table.

### Split-luciferase complementation assay

The coding sequences of *OsAAI1* and *OsMADS25* were inserted into the N-terminal (nLUC) and C-terminal (cLUC) portions of firefly luciferase (LUC) respectively. Split-luciferase complementation assay was performed according to the method described previously[55] by using the resulting plasmids nLUC-OsAAI1 and cLUC-OsMADS25. The primers used for the SLC assay are listed in S2Table.

### Yeast-one-hybrid (Y1H) assays

The full-length CDS of *OsMADS25* was cloned into the pGADT7 vector, while the fragment promoters were cloned into the pHISi vector. The transformation was conducted with the yeast Y1HGold strain. Positive transformants were selected in the medium of SD/-Trp/-Leu containing with or without 50 mM 3-AT (3-amino-1,2,4-triazole).

### Dual-luciferase reporter (DLR) assays

Rice protoplasts were co-transformed with the 35S::OsMADS25 and 35S::OsAAI1 and pGreen II 0800-*Promoter*:LUC using the polyethylene glycol (PEG)-mediated methods, and incubated at 25℃ for 16-h. The firefly luciferase was captured with the luminometer (TANON Chemiluminescent Imaging system), while the LUC activities were determined with a commercial DLR analysis system (DL101-01, Vazyme, Nanjing, China), and the activity was represented by LUC/REN ratios.

### Electrophoretic mobility shift assays (EMSA)

The Electrophoretic mobility shift assays (EMSA) was carried out with reference to Xu et al.[54].

### Paraffin sections

Young roots of rice grown for 5 days were placed in 50% FAA fixative (87 ml ethanol, 10 ml formaldehyde, 3 ml acetic acid) and fixed at 4°C for 48h. The samples were washed three times with phosphate buffer for 20 min each time, dehydrated through 50% ethanol for 2h each time, and the samples were passed through a transparent (85% ethanol for 2h, 95% ethanol (with a small amount of senna) for 2h, anhydrous ethanol for 2h, and 3/4 anhydrous ethanol with 1/4 xylene for 1.5h, 1/2 anhydrous ethanol with 1/2 xylene for 1.5h, 1/4 anhydrous ethanol with 3/4 xylene for 1.5h, and xylene for 2h) before being transferred to paraffin wax containing 1/2 xylene, and placed in a 42°C oven overnight. The material was transferred to a 60°C oven for 4 h, then the paraffin wax was replaced once and kept in a 60°C oven for 2 h. The paraffin wax was replaced again and kept in a 60°C oven for 4 h; and embedded in paraffin. The embedded paraffin blocks were trimmed and sliced with a Reichert HistoSTAT 820 paraffin microtome to a thickness of 5 μm. A drop of water was placed on the slide and the paraffin slices were placed on a drying table and stored in a 40°C oven. The prepared paraffin sections were deparaffinized with xylene three times for 20 min each time, and after deparaffinization, they were stained with 1% solid green. The slides were sealed with neutral adhesive and stored in an oven at 40°C. The slides were observed under a Leica DM6 microscope and photographed.

### Rhizotron experiment

A rhizotron experiment was performed as described previously[56], with slight modifications. Rhizotrons (polymethyl methacrylate containers) with a volume of 1800cm^3^ (20cm ×30cm ×3cm) were used in this study. The rhizotrons were filled with a mixture of vermiculite and nutritious soil (1:1), and germinated seeds were grown in the rhizotrons (1 seed/rhizotron). The inclination angle of rhizotron was adjusted to ∼45° in a greenhouse at 30°C. RAS under drought treatment: germinated seeds were grown in the rhizotrons (1 seed/rhizotron) and the inclination angle of rhizotron was adjusted to ∼45°in a greenhouse at 30°C for one month, and then water cut-off treatment was carried out for 2 months. Images from the rhizotron experiment were analyzed using the EZ-RHIZO/software analysis platform to get an overview of rice RSA[57].

### Physiological Measurements

Plant leaves subjected to drought stress (natural drought, 10% PEG6000, and 20% PEG6000 treatment) for a period of 14 days were utilized for physiological index measurement, with plants under normal conditions serving as the control group. The determination of total chlorophyll content followed the established protocol[58]. MDA content was determined as previously described[59]. Free proline content was measured using the reported method[60]. CAT, APX, GPX activity was measured according to the method as described previously[61,62].

## ACKNOWLEDGEMENTS

This work was supported by the Guizhou Provincial Science and Technology Projects (QKHJC-zk (2021) General 126); China National Tobacco Corporation Guangxi Zhuang Autonomous Region Company (Systematic investigation and establishment of green prevention and control system of tobacco diseases, pests and weeds in Guangxi) (202145000024006).

## AUTHOR CONTRIBUTIONS

Conceptualization: Ning Xu, Haimin Liao, Meng Jiang. Data curation: Jianmin Man. Funding acquisition: Ning Xu, Haimin Liao. Investigation: Ning Xu, Rui Luo, Qing Long. Methodology: Qing Long, Jiaxi Yin. Supervision: Haimin Liao, Meng Jiang. Writing – original draft: Ning Xu. Writing – review & editing: Haimin Liao, Meng Jiang.

## Supporting information

**S1 Table. QRT-PCR primers for IAA-related genes**

**S2 Table. Primer sequences for Yeast two-hybrid assays (Y2H), BiFC and Split-luciferase complementation assay**

**S1 Fig. Paraffin sections of wild-type and transgenic lines.**

Paraffin sections cut transversely(A) and longitudinally(B) of wild-type and transgenic lines were grown normally for 5 days, scale bar = 100μm.

**S2 Fig. RSA of wild type and transgenic lines grown for 30 and 90 days in rhizotrons.**

RSA of wild type and transgenic lines grown for 30 and 90 days in rhizotrons filled with vermiculite and nutritious soil (1:1), scale bar = 6cm, overview of RSA by EZ-RHIZO analysis of images.

**S3 Fig. Phenotypic and statistical analyses of wild-type and transgenic lines treated with 0, 1, 10nM IAA for 7 days.**

Phenology(A), bar = 2cm, plant height(B), primary root length(C), lateral root number(D), lateral root density(E), total root length(F) and adventitious root number(G) of wild type and transgenic lines treated with 0, 0.1, 1, 10nM IAA for 7 days. (Asterisks indicate a statistically significant difference compared with ZH11. \*\**P<0.01*, \*\*\**P <0.001*, \*\*\*\**P<0.0001*; One-way ANOVA).

**S4 Fig Phenotypic of wild-type and transgenic lines treated with 0, 1, 10nM IAA for 14 days.**

Subterranean(A) and above-ground(B) phenotypes of wild-type and transgenic lines grown to 14 days under 0, 0.1, 1, and 10nM IAA treatments, scale bar = 1.5cm.

**S5 Fig. Staining of lateral root primordia of wild-type and transgenic strains grown for 4 days under 0, 0.1, 1, and 10nM IAA treatments.**

Staining of lateral root primordia of wild-type and transgenic strains grown for 4 days under 0, 0.1, 1, and 10nM IAA treatments, scale bar = 0.5cm(A). Statistics on the number of lateral root primordia in test (A)(B). (Asterisks indicate a statistically significant difference compared with ZH11. \*\**P<0.01*, \*\*\**P <0.001*, \*\*\*\**P<0.0001*; One-way ANOVA).

**S6 Fig. Phenotypic and statistical analyses of wild-type and transgenic lines treated with 0, 5nM, 5μM TIBA for 7 days.**

Phenology(A), bar = 2cm, plant height(B), primary root length(C), lateral root number(D), lateral root density(E), total root length(F) and adventitious root number(G) of wild type and transgenic lines treated with 0, 5nM, 5μM TIBA for 7 days. (Asterisks indicate a statistically significant difference compared with ZH11. \*\**P<0.01*, \*\*\**P <0.001*, \*\*\*\**P<0.0001*; One-way ANOVA).

**S7 Fig. Phenology of wild type and transgenic lines treated with 0, 5nM, 5μM TIBA for 14 days, scale bar = 2.5cm.**

**S8 Fig. RSA of wild type and transgenic lines grown for 30 days in rhizotrons with water-off drought treatment.**

RSA of wild type and transgenic lines grown for 30 days in rhizotrons filled with vermiculite and nutritious soil (1:1) supplemented with water-off drought treatment, overview of RSA by EZ-RHIZO analysis of image, bar = 5cm (A). Total root number(B), number of links(C), average length(D), average diameter(E), average projected area(F), average surface area(G) of wild-type and transgenic lines grown with water-off drought treatment for one month in rhizotrons. (Asterisks indicate a statistically significant difference compared with ZH11. \*\**P<0.01*, \*\*\**P<0.001*, \*\*\*\**P<0.0001*; One-way ANOVA).

**S9 Fig. Measurement of physiological indices in wild-type and transgenic lines.** Chlorophyll content(A), proline content(B), MDA content(C), CAT activity(D), APX activity(E), and GPX activity(F) of wild-type and transgenic lines after 14 days of normal, 10% PEG, 10% PEG and 10nM IAA treatments. Chlorophyll content(G), proline content(H), MDA content(I), CAT activity(J), APX activity(K), and GPX activity(L) of ZH11 and *osmads25* after 14 days of 10% PEG, 10% PEG and 10nM IAA treatments. (Asterisks indicate a statistically significant difference compared with ZH11. \*\**P<0.01*, \*\*\**P <0.001*, \*\*\*\**P<0.0001*; One-way ANOVA).

